# Leakage beyond the primary infarction: A temporal analysis of cerebrovascular dysregulation at sites of hippocampal secondary neurodegeneration following cortical photothrombotic stroke

**DOI:** 10.1101/2023.04.24.538047

**Authors:** Rebecca J. Hood, Sonia Sanchez-Bezanilla, Daniel J. Beard, Ruslan Rust, Renée J. Turner, Shannon M. Stuckey, Lyndsey E. Collins-Praino, Frederick R. Walker, Michael Nilsson, Lin Kooi Ong

## Abstract

We have previously demonstrated that a cortical stroke causes persistent impairment of hippocampal-dependent cognitive tasks concomitant with secondary neurodegenerative processes such as amyloid-β accumulation in the hippocampus, a region remote from the primary infarct. Interestingly, there is emerging evidence suggesting that deposition of amyloid-β around cerebral vessels may lead to cerebrovascular structural changes, neurovascular dysfunction, and disruption of blood-brain barrier integrity. However, there is limited knowledge about the temporal changes of hippocampal cerebrovasculature after cortical stroke. In the current study, we aimed to characterise the spatiotemporal cerebrovascular changes after cortical stroke. This was done using the photothrombotic stroke model targeting the motor and somatosensory cortices of mice. Cerebrovascular morphology as well as the colocalization of amyloid-β with vasculature and blood-brain-barrier integrity were assessed in the cortex and hippocampal regions at 7, 28 and 84 days post-stroke. Our findings showed transient cerebrovascular remodelling in the peri-infarct area up to 28 days post-stroke. Importantly, the cerebrovascular changes were extended beyond the peri-infarct region to the ipsilateral hippocampus and were sustained out to 84 days post-stroke. When investigating vessel diameter, we showed a decrease at 84 days in the peri-infarct and CA1 regions that was exacerbated in vessels with amyloid-β deposition. Lastly, we showed sustained vascular leakage in the peri-infarct and ipsilateral hippocampus, indicative of a compromised blood-brain-barrier. Our findings indicate that hippocampal vasculature may represent an important therapeutic target to mitigate the progression of post-stroke cognitive impairment.

## Introduction

Over the recent years, our understanding of secondary neurodegeneration (SND) after stroke has expanded based on pre-clinical and clinical findings. Post-stroke SND involves progressive loss of brain tissue at remote regions connected to the area damaged by the initial infarction that develops within days after the initial stroke and persists for months to years (1, 2). Post-stroke SND has been consistently observed in remote regions such as the ipsilateral thalamus, substantia nigra and white matter tracts in both rodent and human studies (3, 4, 5, 6). The key features of SND include neuronal loss, neuroinflammation and accumulation of neurotoxic proteins (7, 8). This observation is of particular interest as SND shares common pathophysiological features with other neurodegenerative conditions, including tau hyperphosphorylation and amyloid-β deposition (8), suggesting a link between SND and the onset of dementia. Indeed, a previous history of stroke is a major risk factor for development of dementia, with approximately 30% of stroke survivors going on to develop dementia (9, 10).

Our research team has been particularly interested in hippocampal SND after cortical stroke as the hippocampus is known to play a critical role in cognitive function. We recently identified that cortical photothrombotic stroke induces persistent cognitive impairment in various hippocampal-dependent tasks such as associative memory, and cognitive flexibility (11, 12). While the primary infarct was confined to the motor and somatosensory cortices, we demonstrated that the neuropathology changes, including neuronal loss, activation of resident inflammatory cells such as microglia and astrocytes, and increased accumulation of neurotoxic proteins were extended to the ipsilateral hippocampus (11, 12). Interestingly, we showed that the spatial distribution of amyloid-β shifted over time, with a scattered distribution of amyloid-β in the brain parenchyma between 7 to 28 days post-stroke, followed by deposition of amyloid-β around cerebral vessels at 84 days post-stroke (11). While previous studies in Alzheimer’s disease models have shown that deposition of amyloid-β around cerebral vessels may lead to cerebrovascular structural changes, neurovascular dysfunction, and disruption of blood-brain barrier (BBB) integrity (13, 14), little is known about the temporal changes of hippocampal cerebrovasculature after cortical photothrombotic stroke.

The neurovascular unit (NVU; including neurons, astrocytes, microglia, pericytes and endothelial cells) plays an important role in regulating and maintaining cerebral homeostasis, cerebral blood flow and inducing the BBB (15). Recent research has highlighted the importance of the NVU in acute, sub-acute and chronic stroke pathophysiology and led stroke researchers to reconsider the important role of the NVU in acute stroke therapy and recovery

(16). BBB permeability has been observed in humans to be increased from as early as <6 hours to more than 30 days post-stroke and is associated with worse clinical outcomes (17). Despite the potentially detrimental effect of BBB breakdown on stroke outcome in the acute phase of stroke, including hemorrhagic transformation and vasogenic oedema (18), BBB breakdown is a necessary step for the initiation of angiogenesis within the lesion, which is one of the most important processes for functional recovery after stroke (19). In the sub-acute phase of stroke (1-3 weeks), angiogenesis begins resulting in newly formed vessels that are leaky in surrounding and remote regions of the initial stroke injury, and contribute further to BBB permeability (20). However, BBB permeability in these new vessels gradually decreases after 6 weeks, with the result being larger mature vessels that can restore cerebral blood flow to the previously ischemic areas, paving the way for neurogenesis and functional recovery (21). Several studies have documented the importance of angiogenesis in restoring blood flow within the lesion and promoting functional recovery in both animal models (22) and in human patients with stroke (23). However, little is known about temporal changes to angiogenesis, the angioarchitecture and BBB permeability in more distal regions, such as the hippocampus, after cortical stroke, and whether these changes are associated with SND processes.

The aim of the current study was to investigate the impact of cortical photothrombotic stroke on the cerebrovascular morphological changes within the primary cortical infarction site and ipsilateral hippocampal regions at 7, 28 and 84 days post-stroke. Furthermore, we assessed co-localisation of amyloid-β with vasculature, as well as BBB integrity and major tight junction protein and matrix metallopeptidase.

## Materials and Methods

The data that supports the study findings are available from the corresponding author upon reasonable request. See (11) for detailed protocols.

### Experimental Design

All animal experiments were approved by the University of Newcastle Animal Care and Ethics Committee (A-2013-340) and conducted in accordance with the New South Wales Animal Research Act and the Australian Code of Practice for the use of animals for scientific purposes. This study represents an extension of a previous study (11). Therefore, the materials from the same mouse cohort were used to obtain the data featured in this paper. This is in line with the aim to improve the ethical use of animals in testing according to the 3R principles (24).

Briefly, C57BL/6 mice (male, 10 weeks old, n=85) were obtained from the Animal Services Unit (University of Newcastle, Australia). All animals underwent either cortical photothrombosis (n=65) or sham surgery (n=20). Brains were collected at 7, 28 and 84 days post-stroke and used for histological (Sham, n=10; 7 days, n=9; 28 days, n=9; 84 days, n=11) or protein analyses (Sham, n=8; 7 days; n=8, 28 days, n=8; 84 days, n=11). All the experimental groups were randomised, and all outcome analysis performed in a blinded manner in accordance with the ARRIVE guidelines (25).

It should be noted that the sham group consisted of mice subjected to the sham surgical procedure and euthanased at 84 days post sham operation. Based on our preliminary data (26), there were no significant changes in the number of neurons and microglia status between sham groups euthanased between 1 and 182 days post sham operation. Further, previous studies have shown using MRI neuroimaging that no significant changes occur in the brain volume of sham operated animals over this period (27). Therefore, in this study we have only included the sham group corresponding to the 84 day time point.

### Photothrombotic Occlusion

Photothrombotic occlusion (and sham surgery) was performed as previously described (11, 28, 29, 30). Briefly, following isoflurane anaesthesia (2%), 200 µL of Rose Bengal (10mg/mL in sterile saline; Sigma Aldrich, USA) or vehicle control was injected intraperitoneally and permitted to circulate for 8 min. In parallel to circulation, an incision was made in the scalp to expose the skull, which was then illuminated by a 4.5 mm diameter cold light source for 15 min, positioned at 2.2 mm lateral to bregma over the left motor and somatosensory cortices.

### Tissue Processing

Mice were euthanised at 7, 28, or 84 days post-stroke and brains were prepared for histological or protein analyses as previously described (11, 12, 31). For histological analysis, mice were deeply anaesthetised using sodium pentobarbitol (I.P.) and transcardially perfused with 10 mL ice-cold saline (0.9%) followed by 40 mL ice-cold paraformaldehyde (4%, pH 7.4). Brains were collected and fixed for a further 4 hours in 4% paraformaldehyde before cryoprotection in 12.5% sucrose in 0.1 M PBS. Brains were coronally sectioned at 30 µm on a Freezing microtome (Leica, Australia). For Western blotting, mice were deeply anaesthetised using pentobarbitol (I.P.) and transcardially perfused with ice-cold 0.1% diethyloyrocarbonate in 0.9% saline for at least 3 min, or until the blood ran clear. Brains were rapidly collected and frozen in −80°C isopentane. Coronal sections were sectioned at 200 µm using a cryostat (Leica, Australia) set at −20 °C.

### Immunoflourescence

Free floating fixed sections (30 µm) were labelled for lectin, PDGFRβ, amyloid-β and IgG using previously described methods (11, 32, 33). Briefly, sections were rinsed and non-specific binding blocked using 3% bovine serum albumin. Primary antibodies (PDGFRβ, amyloid-β and IgG) were applied and sections incubated overnight at 4 °C, followed by incubation in the corresponding secondary antibodies at 25 °C for 2 hours. Lectin was added in parallel to secondary antibody incubation. See Table 1 for antibodies and concentrations. Brain sections were washed with PBS in between each incubation step. Sections were then mounted and cover slipped.

**Table 1:** List of antibodies used for western blot and immunofluorescence analyses.

| Target | Sources of antibodies | Application | Dilution |
| --- | --- | --- | --- |
| **Amyloid-β** | Biologend, anti-Amyloid-β (6E10), # SIG-39320 | IF | 1:1000 |
| **Claudin 5** | Invitrogen, Claudin 5 Polyclonal Antibody, #34-1600 | WB | 1:1000 |
| **Lectin** | Vecton Laboratories, DyLight 649 Lycopersicon esculentum (Tomato) lectin #DL-1178 | IF | 1:1000 |
| **MMP9** | Abcam, Anti-MMP9 antibody, #ab38898 | WB | 1:1000 |
| **PDGFRβ** | Abcam, Recombinant Anti-PDGFR-alpha + PDGFR-beta antibody [Y92] C-terminal, #ab32570 | IF | 1:1000 |
| **β-actin** | Sigma-Aldrich, Monoclonal Anti-β-actin-HRP antibody, A3854 | WB | 1:50000 |
| **Rabbit IgG** | Biorad, Anti-Rabbit-HRP antibody, #170-6515 | WB | 1:7500 |
|  | ThermoFisher Scientific, anti-Rabbit IgG (H+L) Highly Cross-Adsorbed Secondary Antibody, Alexa Fluor 488, #A21206 | IF | 1:400 |
| **Mouse IgG** | Biorad, Anti-Mouse-HRP antibody, #170-6516 | WB | 1:10000 |
|  | ThermoFisher Scientific, anti-Mouse IgG (H+L) Highly Cross-Adsorbed Secondary Antibody, Alexa Fluor 594, #A21203 | IF | 1:400 |

WB, western blot; IF, immunofluorescence

### Image Quantification

High resolution images were acquired using a Leica TCS SO8 confocal microscope with Leica HC PLC APO 20x/0.70 and 10x/0.40 objectives for the peri-infarct and hippocampal regions, respectively. Z-stacks (30µm with a step size of 1µm) were taken for each region of interest with imaging parameters maintained throughout imaging sessions (laser power, resolution and gain).

Quantitative analysis was performed on the ipsilateral motor (M1) and somatosensory (S1) cortices (within the peri-infarct territory; Bregma 0.0 mm; Figure 1A), as well as the CA1 and DG sub-regions of the ipsilateral hippocampus (Bregma –1.5) (Figure 2A). Vascular images were pre-processed and analyzed as previously established with an automated ImageJ (Fiji) script (34, 35). In brief, images were duplicated, transformed to 8-bit, and processed with a median filter to remove noise. Images were then binarized, allocating vascular signal the value 255 (non-zero-pixel) and the background signal value 0 (zero-pixel) (see Supplementary Figure 1). The following parameters were calculated based on the binarized images:

1. *Area fraction (%) of the blood vessels or pericytes*: Area fraction calculation measures the percentage of pixels with non-zero pixels to all pixels.
2. *Vessel length per mm^2^*: Vessel segment length and was assessed by skeletonizing the binary image. This allowed us to tag all pixels in a skeletonised image and then measure their length (mm). The length was normalized per mm^2^ of analyzed brain tissue. Unit: mm/mm^2^
3. *Number of vascular branches per mm^2^*: The number of branches was assessed by skeletonizing the binary image. This allowed us to identify all its branches. The number of branches was normalized per mm^2^ of analyzed brain tissue. Unit: 1/mm^2^
4. *Average vessel diameter:* Fifteen random vessels were sampled from each binary image and manually traced for average vessel diameter. Colocalization of lectin and amyloid-β labels was used to visualize where both labels overlap for the measurement of amyloid-β positive or negative vessel diameter at 84 days post stroke. Unit: µm
5. PDGFRβ+ pericyte coverage of the vasculature *(%):* The pericyte coverage is quantified by dividing the PDGFRβ area fraction signal from lectin area fraction signal.

**Figure 1:**
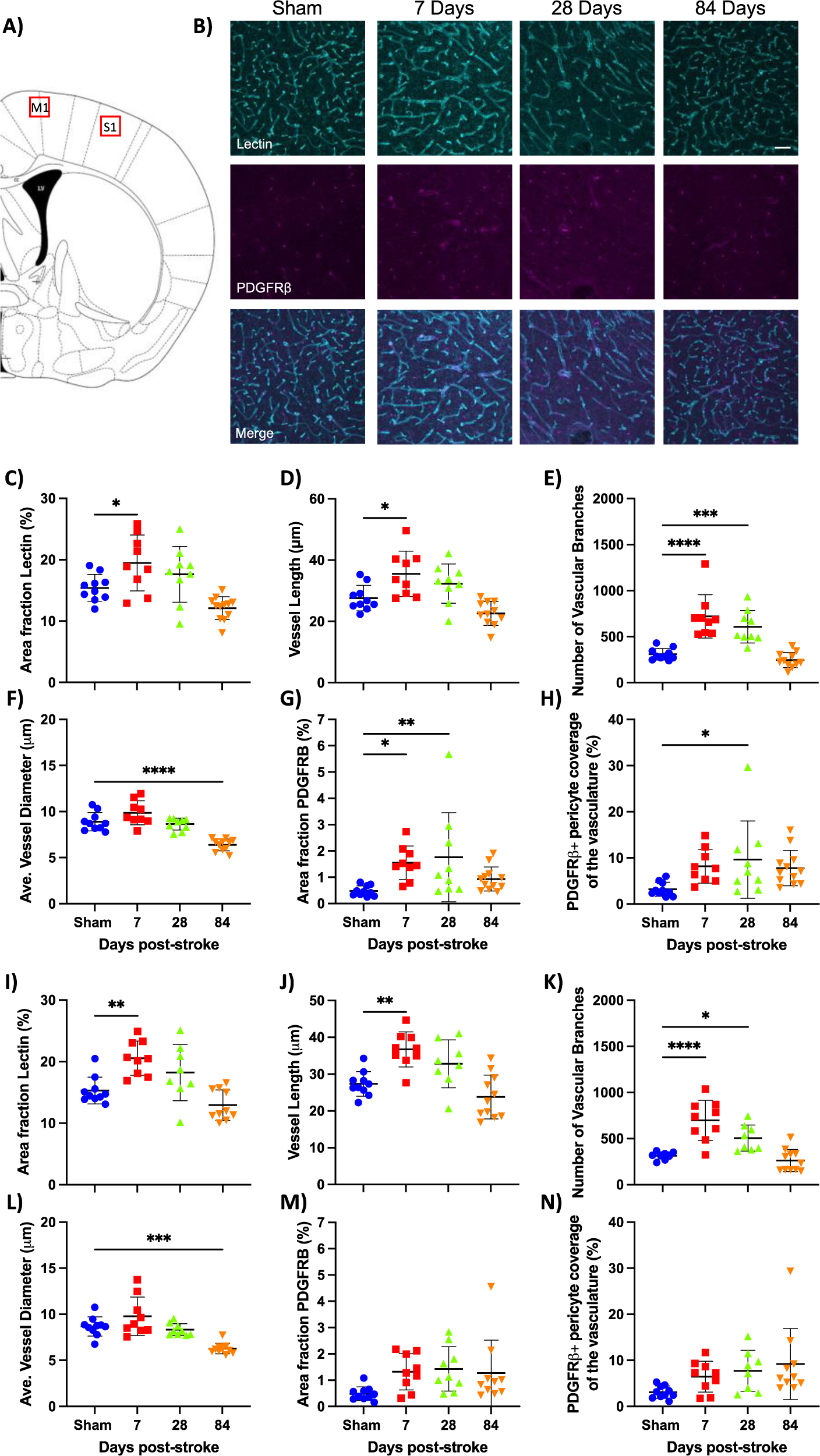
Temporal changes in the cerebrovascular morphology of the peri-infarct area after cortical photothrombotic stroke. A) Schematic showing the location of image analysis in the motor (M1) and sensory (S1) cortices. B) Representative immunofluorescence labelling for lectin (cyan) and PDGFRβ (magenta) up to 84 days post-stroke. Changes to lectin and PDGFRβ over time in the M1 (C-H) and S1 regions (I-N). Scale bar = 50 µm. Mean ± SD (Two-way ANOVA with Tukey’s multiple comparisons test). *p<0.05. **p<0.01. ***p<0.001.

**Figure 2:**
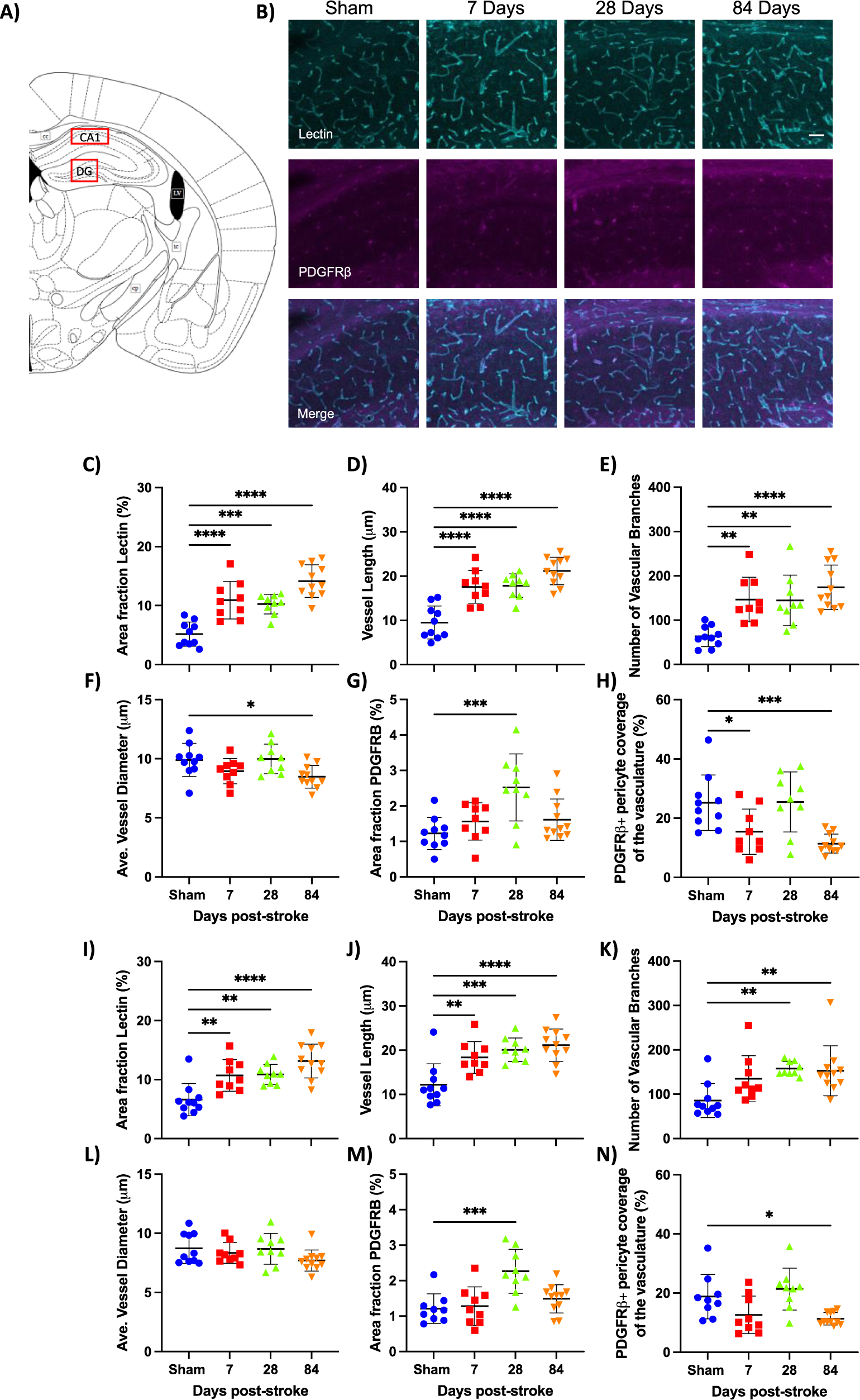
Temporal changes in the cerebrovascular morphology of hippocampal sub-regions after cortical photothrombotic stroke. A) Schematic showing the location of image analysis in the CA1 and dentate gyrus (DG) of the hippocampus. B) Representative immunofluorescence labelling for lectin (cyan) and PDGFRβ (magenta) up to 84 days post-stroke. Changes to lectin and PDGFRB over time in the CA1 (C-H) and DG sub-regions (I-N). Scale bar = 50 µm. Mean ± SD (Two-way ANOVA with Tukey’s multiple comparisons test). *p<0.05. **p<0.01. ***p<0.001.

For IgG labelling, we performed thresholding analysis (as previously described (11, 12, 36), and investigated the percentage of thresholded material across different pixel intensity thresholds (0-255).

### Protein extraction and Western Blotting

Protein homogenates were obtained from the peri-infarct (bregma +1.0 to –1.0 mm) and the ipsilateral hippocampus (Bregma −1.2 to –2.5 mm) for Western blot analysis as previously described (11, 12, 28, 31). Briefly, tissue was collected using a 1 mm tissue punch and sonicated in 300µL lysis buffer (50 mM TRIS buffer pH 7.4, 1 mM EDTA, 1 mM DTT, 80 μM ammonium molybdate, 1 mM sodium pyrophosphate, 1 mM sodium vanadate, 5 mM b-glycerolphosphate, 1 protease inhibitor cocktail tablet, 1 phosphatase inhibitor cocktail tablet, final concentration) and centrifuged at 14000 G for 20 min at 4 °C. The supernatant was collected and protein concentration was quantified using a Pierce BCA protein assay kit (Thermo Fisher Scientific, USA) according to manufacturer instructions. Sample buffer (2% SDS, 50 mM Tris, 10% glycerol, 1% DTT, 0.1% bromophenol blue, pH 6.8) was then added to samples. The samples (15 μg of total protein) were then loaded into Biorad Criterion TGX Stain-Free 4–20% gels for electrophoresis. Proteins were transferred from the gels to PVDF membranes, washed in Tris-buffered saline with tween (TBST; 150 mM NaCl, 10 mM Tris, 0.075% Tween-20, pH 7.5) and incubated in 5% skim milk powder in TBST for 1 hour at room temperature. Membranes were incubated with primary antibodies (Claudin 5 and MMP9) at 4 °C overnight, followed by and secondary antibody for 1 hour at room temperature (see Table 1 for antibody concentration). Membranes were washed in TBST between each incubation step. Membranes were visualized on an Amersham Imager 600 using Luminata Classico or Luminata Forte western blotting detection reagents. The density of the bands was measured using Amersham Imager 600 analysis software. See Supplementary Materials for raw western blots.

### Statistical Analysis

Data were analysed using GraphPad Prism v7.02. The primary outcome measure was the difference between sham and the experimental groups (7, 28 and 84 days post-stroke). Data were tested for normality and then analysed using a one-way analysis of variance (ANOVA) and corrected for multiple comparisons using a Dunnett’s post-hoc analysis. Average vessel diameter of amyloid-β positive vs amyloid-β negative at 84 days post stroke were analysed using a paired t-test. All data are presented as mean ± SD. Statistical significance was accepted at p<0.05.

## Results

### Cortical photothrombotic stroke causes transient cerebrovascular remodelling in the peri-infarct area

#### Lectin

In both peri-infarct regions (M1 and S1), mice exhibited a transient increase in lectin staining, vessel length and number of branches (Figure 1B-E, I-K). Post-hoc testing showed that the area fraction of lectin (%) was significantly higher in both regions at day 7 (M1, p=0.0344; S1, p=0.0019) before returning to sham levels at 28 days. They also exhibited an increase in vessel length (M1, p=0.0103; S1, p=0.0013) and number of junctions (M1, p<0.0001; S1, p<0.0001) at 7 days, the latter of which extended out to 28 days post-stroke (M1, p=0.0004; S1, p=0.0199) before returning to sham levels at 84 days. In both regions, average vessel diameter was significantly decreased from sham levels at 84 days post-stroke (M1, p<0.0001; S1, p=0.0002), but not at any of the other time points (Figure 1F, L). See Table 3 for a summary of PI findings.

**Table 2:** Summary of findings in the peri-infarct area.

|  | Motor Cortex (M1) |  |  |  | Sensory Cortex (S1) |  |  |  |
| --- | --- | --- | --- | --- | --- | --- | --- | --- |
|  | 2 way ANOVA | 7 d | 28 d | 84 d | 2 way ANOVA | 7 d | 28 d | 84 d |
| Cerebrovascular Remodelling |  |  |  |  |  |  |  |  |
| Area fraction Lectin | F<sub>(3,36)</sub>=9.239, p=0.0001 | ↑ | NS | NS | F<sub>(3,33)</sub>=11.26, p<0.0001 | ↑↑ | NS | NS |
| Vessel length | F<sub>(3,35)</sub>=10.29, p<0.0001 | ↑ | NS | NS | F<sub>(3,34)</sub>=11.69, p<0.0001 | ↑↑ | NS | NS |
| Number of vascular branches | F<sub>(3,35)</sub>=22.42, p<0.0001 | ↑↑↑↑ | ↑↑↑ | NS | F<sub>(3,34)</sub>=18.83, p<0.0001 | ↑↑↑↑ | ↑ | NS |
| Average vessel diameter | F<sub>(3,35)</sub>=26.08, p<0.0001 | NS | NS | ↓↓↓↓ | F<sub>(3,35)</sub>=14.81, p<0.0001 | NS | NS | ↓↓↓ |
| Area fraction PDGFRβ | F<sub>(3,36)</sub>=4.103, p=0.0103 | ↑ | ↑↑ | NS | F<sub>(3,34)</sub>=2.547, p=0.0722 | - | - | - |
| PDGFRβ+ pericyte coverage of the vasculature | F<sub>(3,36)</sub>=3.183, p=0.0354 | NS | ↑ | NS | F<sub>(3,33)</sub>=2.842, p=0.0528 | - | - | - |
| Vascular Leakage (IgG) |  |  |  |  |  |  |  |  |
| Pixel Intensity 210 |  | ↑ | NS | NS |  | ↑ | NS | NS |
| Pixel Intensity 230 | F<sub>(3,8960)</sub>=586, p<0.0001 | ↑↑↑↑ | ↑↑ | ↑↑ | F<sub>(3,8960)</sub>=632.8, p<0.0001 | ↑↑↑↑ | ↑↑↑ | ↑↑ |
| Pixel Intensity 240 |  | ↑↑↑↑ | ↑↑↑↑ | ↑↑↑↑ |  | ↑↑↑↑ | ↑↑↑↑ | ↑↑↑↑ |
| Pixel Intensity 245 |  | ↑↑↑↑ | ↑↑↑↑ | ↑↑↑↑ |  | ↑↑↑↑ | ↑↑↑↑ | ↑↑↑↑ |
|  | Peri-infarct Area |  |  |  |  |  |  |  |
| BBB Proteins | F Statistic | 7 d | 28 d | 84 d |  |  |  |  |
| Claudin 5 | F<sub>(3,31)</sub>=26.9, p<0.0001 | NS | ↓↓↓↓ | ↓↓↓↓ |  |  |  |  |
| MMP9 | F<sub>(3,31)</sub>=65.62. p<0.0001 | ↑↑↑↑ | ↑ | NS |  |  |  |  |

Arrow=\*, NS=not significant (post-hoc), - ANOVA was not significant therefore no post-hoc comparison.

**Table 3:** Summary of findings in the hippocampus and sub-regions.

|  | CA1 Subregion |  |  |  | Dentate Gyrus (DG) |  |  |  |
| --- | --- | --- | --- | --- | --- | --- | --- | --- |
|  | 2 way ANOVA | 7 d | 28 d | 84 d | 2 way ANOVA | 7 d | 28 d | 84 d |
| **Cerebrovascular Remodelling** |  |  |  |  |  |  |  |  |
| Area fraction Lectin | $F_{(3,35)}=23.01$,<br>$p<0.0001$ | ↑↑↑↑ | ↑↑↑ | ↑↑↑↑ | $F_{(3,35)}=11.70$,<br>$p<0.0001$ | ↑↑ | ↑↑ | ↑↑↑↑ |
| Vessel length | $F_{(3,35)}=22.6$,<br>$p<0.0001$ | ↑↑↑↑ | ↑↑↑↑ | ↑↑↑↑ | $F_{(3,35)}=11.47$,<br>$p<0.0001$ | ↑↑ | ↑↑↑ | ↑↑↑↑ |
| Number of vascular branches | $F_{(3,35)}=10.63$,<br>$p<0.0001$ | ↑↑ | ↑↑ | ↑↑↑↑ | $F_{(3,35)}=5.463$,<br>$p<0.0035$ | NS | ↑↑ | ↑↑ |
| Average vessel diameter | $F_{(3,35)}=3.934$,<br>$p=0.0161$ | NS | NS | ↓ | $F_{(3,35)}=1.945$,<br>$p=0.1404$ | - | - | - |
| Area fraction PDGFRβ | $F_{(3,35)}=6.857$,<br>$p=0.0009$ | NS | ↑↑↑ | NS | $F_{(3,34)}=8.586$,<br>$p=0.0002$ | NS | ↑↑↑ | NS |
| PDGFRβ+ pericyte coverage of the vasculature | $F_{(3,35)}=8.117$,<br>$p=0.0003$ | ↓ | NS | ↓↓↓ | $F_{(3,34)}=6.245$,<br>$p=0.0017$ | NS | NS | ↓ |
| **Vascular Leakage (IgG)** |  |  |  |  |  |  |  |  |
| Pixel Intensity 210 |  | - | - | - |  | - | - | - |
| Pixel Intensity 230 | $F_{(3,8960)}=281.8$,<br>$p<0.0001$ | - | - | - | $F_{(3,8960)}=84.65$,<br>$p<0.0001$ | - | - | - |
| Pixel Intensity 240 |  | ↑↑↑↑ | ↑↑↑↑ | ↑ |  | ↑ | ↑ | NS |
| Pixel Intensity 245 |  | ↑↑↑↑ | ↑↑↑↑ | ↑↑ |  | NS | ↑↑ | NS |
|  | **Hippocampus** |  |  |  |  |  |  |  |
| **BBB Proteins** | F Statistic | 7 d | 28 d | 84 d |  |  |  |  |
| Claudin 5 | $F_{(3,31)}=5.524$, $p=0.0037$ | ↑↑ | NS | NS | | | | |
| MMP9 | $F_{(3,31)}=7.629$, $p=0.0006$ | NS | ↑ | NS | | | | |

Arrow=\*, NS=not significant (post-hoc), - ANOVA was not significant therefore no post-hoc comparison.

#### PDGFRβ

There was also a transient increase in PDGFRβ labelling in the M1 region at 7 (p=0.0354) and 28 days (p=0.0099) post-stroke, but no change in the S1 region (Figure 1G, M). The PDGFRβ+ pericyte coverage of the vasculature was increased in the M1 region at 28 days (p=0.0186) post-stroke, but there were no changes in the S1 region (Figure 1H, N).

#### Amyloid-β

At 84 days post-stroke in both regions, mean vessel diameter was significantly reduced in vessels that were colocalised with amyloid-β (M1, 5.13±0.8 μm; S1, 4.99±0.6 μm) vs those that were not co-localised (M1, 6.8±0.7 μm; S1, 6.5±0.3 μm) p=0.0001 and p<0.0001, respectively (Figure 3A-C).

**Figure 3:**
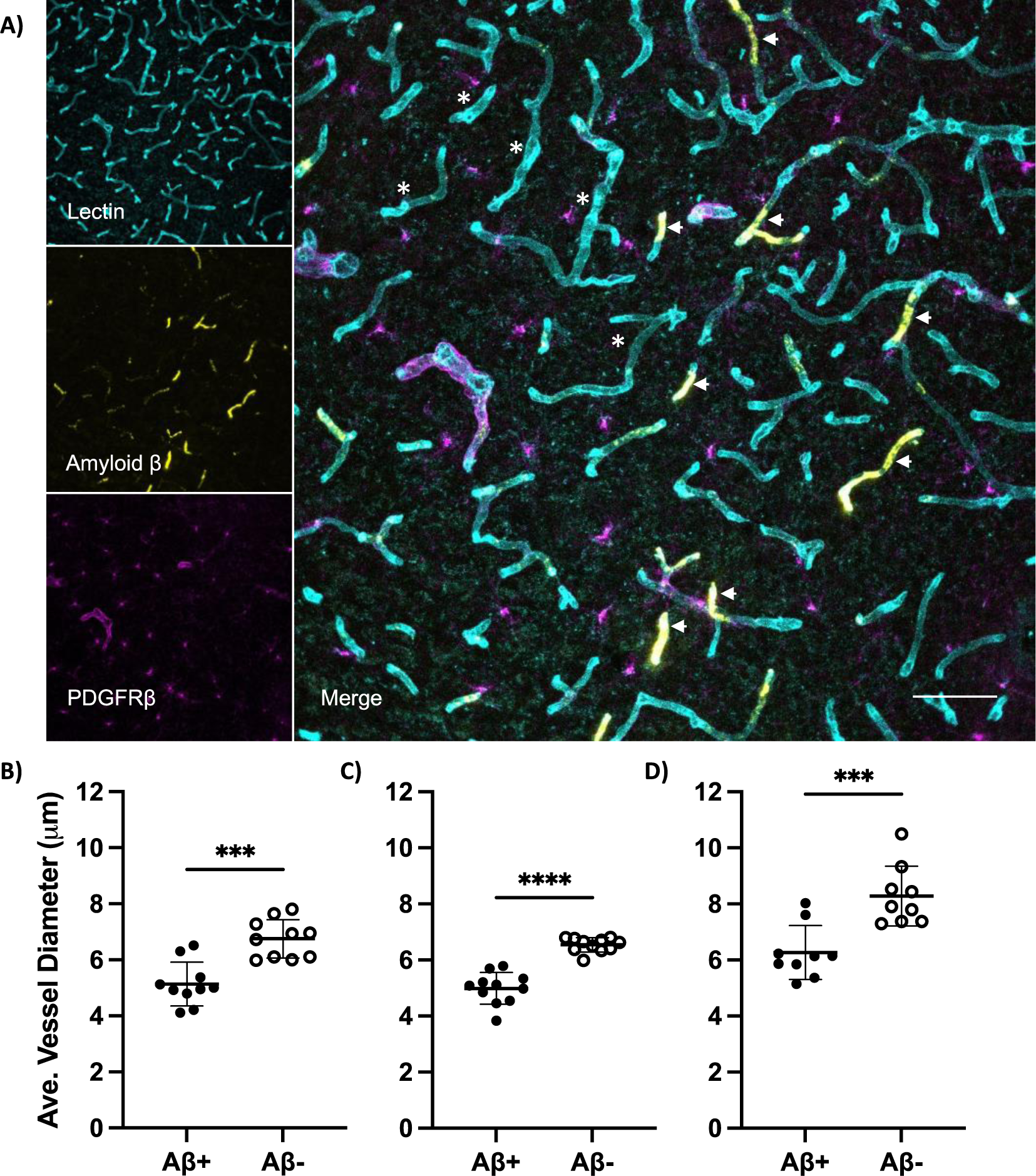
Vessels colocalised with amyloid-β (Aβ) have significantly reduced vessel diameters at 84 days post-stroke. A) Representative immunofluorescence labelling for lectin (cyan), amyloid-β (yellow) and PDGFRβ (magenta). Vessels with amyloid-β deposition (◄, white arrow) showed a decrease in vessel diameter compared to vessels without amyloid-β deposition (*, asterisk). Scale bar = 50 µm. Average vessel diameter in vessels ± Aβ in the B) motor (M1) and C) sensory (S1) cortices, and in the D) CA1 subregion of the hippocampus. ***p<0.001, ****p<0.0001.

### Cortical photothrombotic stroke causes sustained cerebrovascular remodelling in the ipsilateral hippocampus

#### Lectin

In the CA1 subregion of the hippocampus, mice exhibited a sustained increase in lectin staining, vessel length and the number of branches (Figure 2B-E). Post-hoc testing showed increases from sham at all time points across the parameters (p≤0.0017). There was a decrease in average vessel diameter in the CA1 subregion at 84 days post-stroke (p=0.0244), but not at any other time point (Figure 2F). See Table 3 for a summary the findings in the hippocampus and sub-regions.

Similarly, in the DG subregion, mice exhibited changes in lectin staining, vessel length and the number of junctions (Figure 2I-K). Post-hoc testing showed increases from sham across all time points in both lectin staining (p≤0.0037) and vessel length (p≤0.0029). Increases were also observed in the number of branches at 28 (p=0.0032) and 84 days post-stroke (p=0.0041). There was no change in average vessel diameter (Figure 2L).

#### PDGFRβ

Both CA1 and DG subregions also showed a transient increase in PDGFRβ labelling. Post-hoc testing showed elevated levels in both regions at 28 days post-stroke (CA1, p=0.0003 and DG, p=0.0002) (Figure 2G, M). The PDGFRβ+ pericyte coverage of the vasculature also differed over time in both regions. Post-hoc testing showed a decrease at 7 (p=0.0282) and 84 days (p=0.0009) post-stroke in the CA1 subregion and at only 84 days in the DG (p=0.0235) (Figure 2H, N).

#### Amyloid-β

At 84 days post-stroke in the CA1 subregion, mean vessel diameter was significantly reduced in vessels that were colocalised with amyloid-β (6.3±0.96 μm) vs those that were not co-localised (8.3±1.1 μm), p=0.0006 (Figure 3D).

### Cortical photothrombotic stroke increases vascular leakage (IgG) in the peri-infarct area and ipsilateral hippocampus

We measured the IgG immunoreactivity using cumulative threshold analysis. The number of pixels occurring at each of the pixel intensities was determined and the pixel intensity values were ranked ordered 0 to 255 along with the corresponding number of pixels that occurred at each value. As expected, limited immunofluorescence signal was observed within the peri-infarct region and hippocampus in shams. We found a significant increase in IgG immunoreactivity in M1 and S1 region of stroked mice at all time points compared to shams (p<0.0001 at 7 days, 28 days and 84 days). In the CA1 subregion of the ipsilateral hippocampus, we detected a significant increase in IgG immunoreactivity at all time points post-stroke (p<0.0001 at 7 and 28 days, and p=0.0378 at 84 days). In the DG subregion however, we only observed a transient increase in IgG immunoreactivity in the DG at 7 days (p=0.0176) and 28 days (p=0.0139) in stroked mice compared with shams (Figure 4).

**Figure 4:**
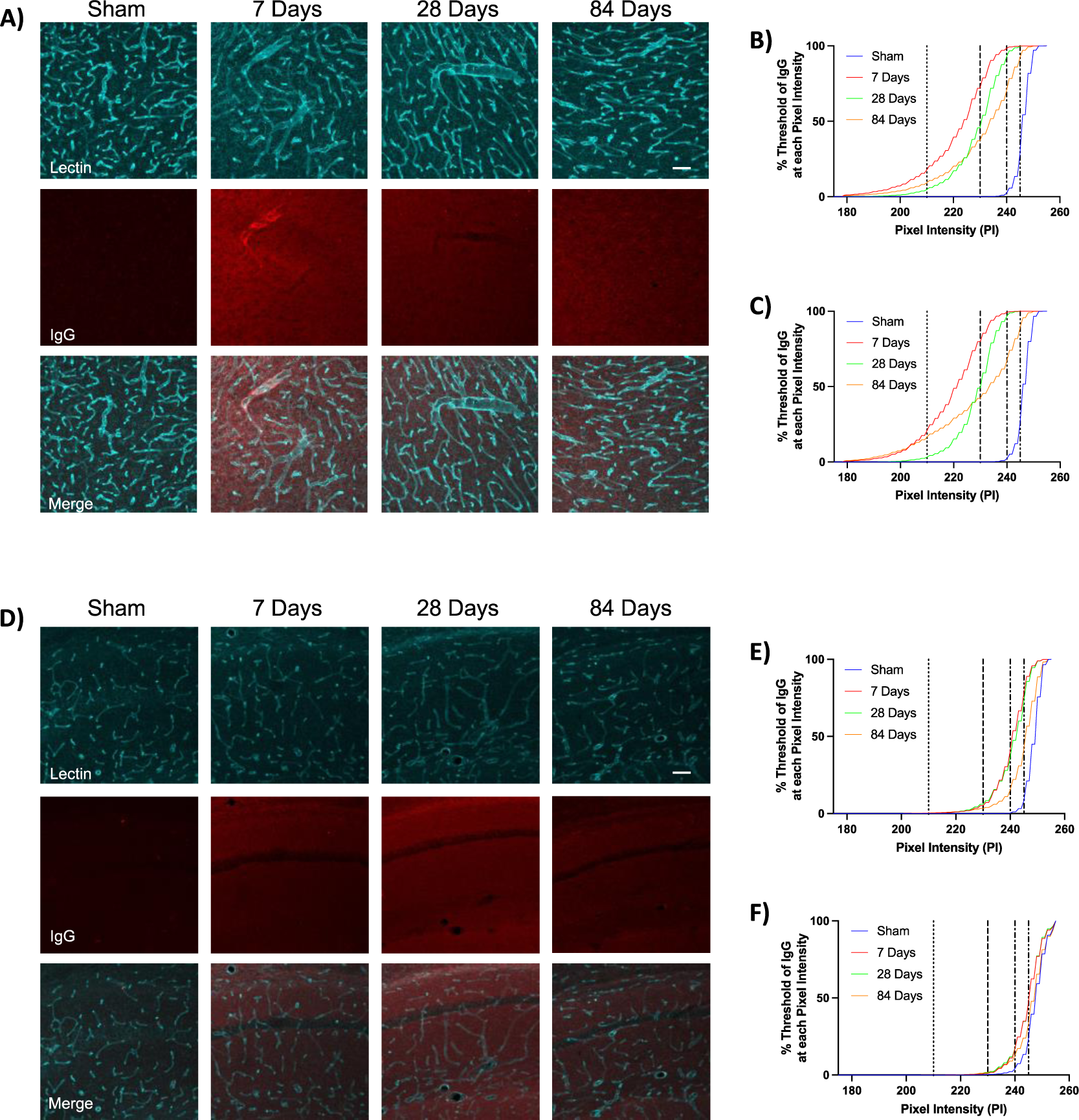
Cortical photothrombotic stroke induces BBB leakage in the peri-infarct area (top panel), and ipsilateral hippocampus (bottom panel). Representative immunofluorescence labelling of lectin (cyan) and IgG (red) up to 84 days post-stroke in the peri-infarct area (A) and hippocampus (D). Scale bar = 50 µm. Cumulative threshold analysis showing the mean percentage of IgG in the M1 (B) and S1 (C) cortices; and CA1 (E) and DG (F) subregions of the hippocampus at different levels of pixel intensity (PI). The dotted lines represent the PI levels used in the analysis for detecting genuine differences in immunoreactive signal.

### Cortical photothrombotic stroke induces transient changes in expression of BBB proteins in the peri-infarct area and ipsilateral hippocampus

To assess temporal changes to the BBB, we investigated Claudin 5 (a tight junction protein expressed in endothelial cells (37)) and MMP9 (an endopeptidase that can degrade the components of extracellular matrix (37)) (Figure 5). In the peri-infarct area, we found effects of time on the fold change of both proteins. Post hoc analysis showed a significantly decreased levels of Claudin 5 at 28 and 84 days post-stroke (both p<0.0001). We also observed a transient increase in MMP9, peaking at 7 days post stroke (p<0.0001) and continuing to be elevated at 28 days (p=0.0174) before returning to sham levels at 84 days. Similarly, in the hippocampus we found effects of time on both Claudin 5 and MMP9. Claudin 5 transiently increased, peaking at 7 days post-stroke (p=0.0013), before returning to sham levels by 28 days. MMP9 also displayed a transient increase, with fold change peaking at 28 days post-stroke (p=0.013) and returning to sham levels by 84 days.

**Figure 5:**
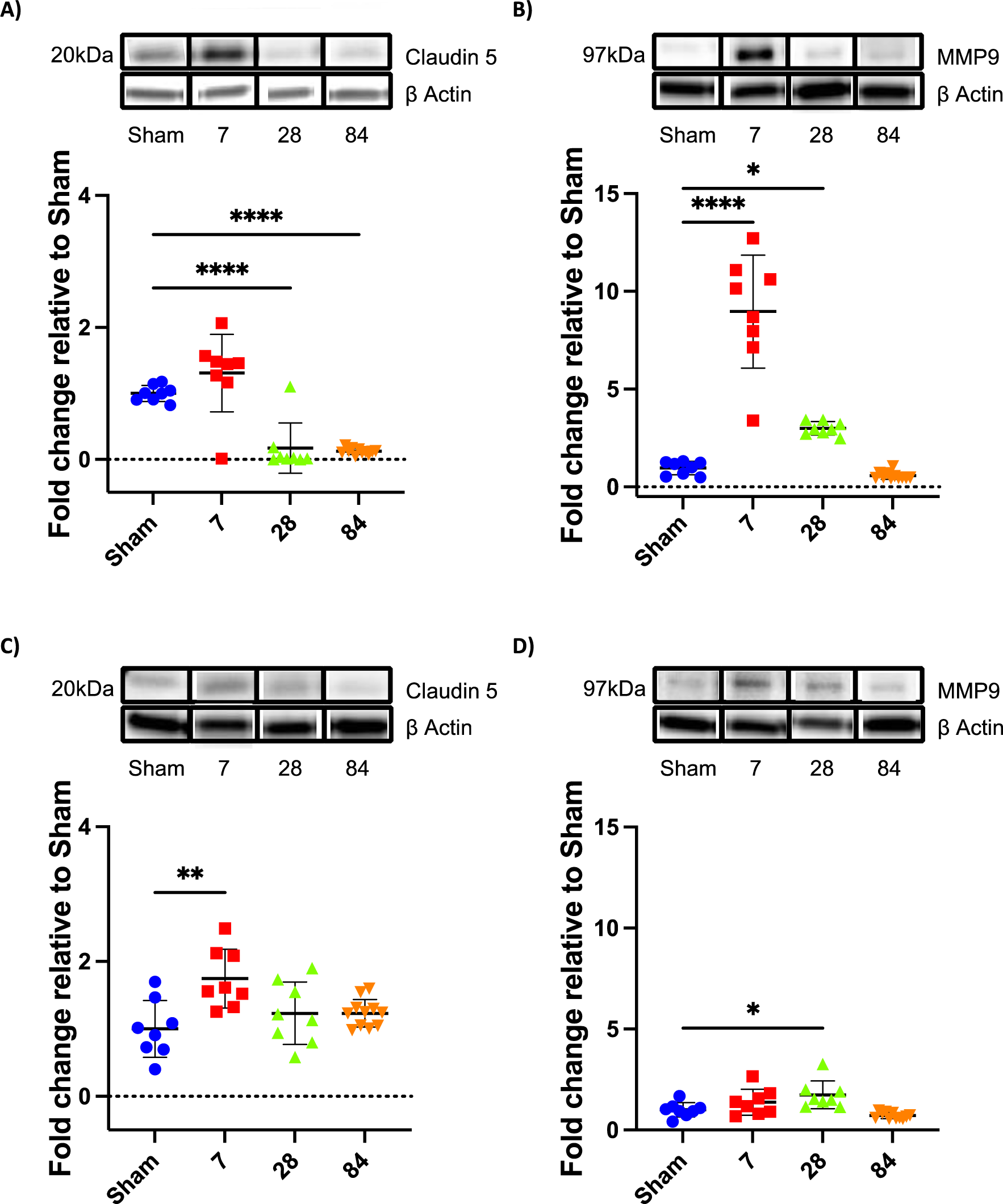
Quantification of Claudin 5 and Matrix metalloproteinase 9 (MMP9) in the peri-infart area ((A) and (B), respectively; top panel) and hippocampus ((C) and (D), respectively; bottom panel) out to 84 days post-cortical photothrombosis. Representative blots of Claudin 5, MMP9 and β-actin. Full western blots can be viewed in Supplementary Figures 2-5. Loading controls were performed by loading equal amounts of total protein and also were normalised to β-actin. Levels were expressed as a fold change (FC) of mean ± SD for each group relative to the mean of the sham group, shown in the graphs for each protein. *p<0.05. **p<0.01. ***p<0.001. ****p<0.0001.

## Discussion

We have previously shown that unilateral cortical photothrombotic stroke causes persistent impairment in associative memory and learning, as well as in cognitive flexibility, up to 84 days post-stroke (Figure 6 (11)). These deficits were associated with concomitant hippocampal neuropathology, including neuronal loss, microglial activation and the accumulation of amyloid-β. Here we extended upon these findings and characterised the spatiotemporal cerebrovascular changes after cortical stroke. We demonstrated for the first time that cortical photothrombosis causes persistent remote hippocampal cerebrovascular dysregulation. Specifically, we showed transient cerebrovascular remodelling (vessels and PDGFRβ+ pericytes) in the peri-infarct areas. Interestingly, the cerebrovascular changes were extended beyond the peri-infarct region to the ipsilateral hippocampus, a site recently documented to undergo SND after cortical stroke, and were sustained out to 84 days post-stroke. When investigating vessel diameter, we showed a decrease at 84 days in the peri infarct areas and CA1 regions that was exacerbated in vessels with amyloid-β deposition. Lastly, we showed sustained vascular leakage in the peri-infarct areas and ipsilateral hippocampus, indicative of a compromised BBB. Collectively, these results suggest that cortical stroke induces remote hippocampal cerebrovascular dysregulation, particularly reduction of vessel diameter, as well as BBB leakage, which may partly contribute to the progression of post-stroke SND and cognitive impairment.

**Figure 6:**
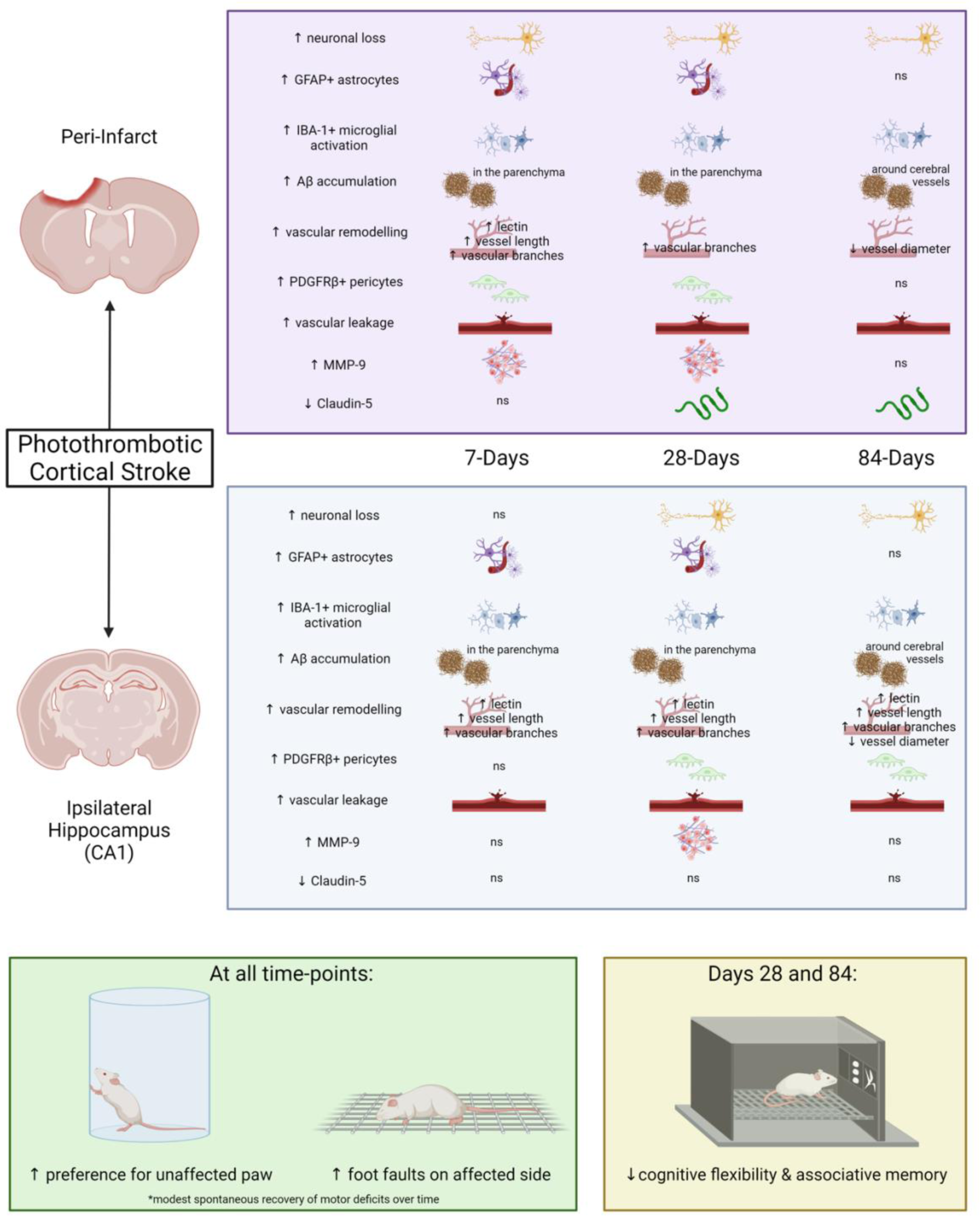
Overview of spatiotemporal changes observed in mice after cortical photothrombotic stroke. Includes findings from Sanchez-Bezanilla *et al.* (2021) (11). Ns= not significant. Figure created with BioRender.com

In this study, we showed an initial increase in vessel density, vessel length and the number of vascular branches within the peri-infarct areas at 7 days post-stroke. However, these vessel changes had returned to sham levels by 28-84 days post-stroke. Similar increases in peri-infarct blood vessel number have been reported in mice after middle cerebral artery occlusion (MCAo) as early as 2-3 days post-stroke (22, 38). Evidence of vascular proliferation out to 4 weeks post-stroke in peri-infarct areas has also been observed in post-mortem tissue samples from stroke patients (39). Further, Krupinski *et al.* (1994) showed that angiogenesis within the penumbra correlated with longer post-stroke survival in a small number of patients (23). Although we did not determine whether the vessels were capable of flow, Morris *et al.* (2022) (40) showed evidence of patency in the peri-infarct region at 2 weeks post stroke using vascular casting with fluorescein isothiocyanate labelled albumin. Suggesting that at least some of the vessels we observed may be capable of flow. Interestingly, the early changes we observed had resolved by 28-84 days post-stroke. This is consistent with findings by Yu *et al.* (2006), who showed no differences in the number of microvessels in the peri-infarct area at 30-, 90- and 165-days post-2h MCAo (41). Although we did not study the mechanisms underlying the cerebrovascular changes, previous studies have reported increases in protein and gene expression of angiogenesis associated markers as early as 1 hour post-stroke compared with the contralateral hemisphere e.g VEGF, flk-1 and angiopoetin-1 (22, 38). Interestingly, the upregulation of many of the markers peaked between 1-7 days and then trended downward out to 21 days. The authors did not look beyond this period, so it is unknown whether protein levels returned to contralateral levels. The observed downregulation of vessels in this study may reflect the downregulation of many of the angiogenesis associated markers observed in the studies. These newly formed vessels would permit increased blood flow and oxygenation to the affected tissue. We hypothesise that the observed angiogenesis and/or vasculogenesis within the motor cortex may have contributed to the modest improvements in motor function observed in the animals used in the study (Figure 6, see (11) for more information). Previous studies have shown similar associations between vascular repair and improvement in motor function after stroke (42).

Further, we observed a transient increase in PDGFRβ staining at 7 and 28 days in the M1 region of the peri-infarct area. A similar increase in PDGFRβ labelling was observed by Fernandez-Klett *et al*. (2013) in the lesion site 5 days post-MCAo, remaining elevated out to 28 days (43). The authors found an increase in proliferation of PDGFRβ+ and CD13+ cells peaking at 3 days but remaining elevated out to 28 days post-stroke. A similar pattern of distribution was observed by the authors in 16 post-mortem tissue samples in the acute to chronic stages post-stroke. The significance of these findings is that pericytes are important in new vessel formation as well as functional recovery (44). Both of which were observed in these animals (11).

There is an increasing body of evidence implicating the hippocampus as a site of SND after stroke (11, 12, 31). In this study, we found persistent increases in vessel density, length and the number of vascular branches out to 84 days post-stroke in the CA1 and DG hippocampal subregions, despite both subregions being remote from the primary infarction. A transient increase in PDGFRβ staining was also observed in the hippocampal subregions, peaking at 28 days post-stroke. Increases in microvascular density have been reported in other areas associated with SND (1), including the ipsilateral thalamus and substantia nigra (45). Hayward *et al*. (2011) found evidence of angiogenesis coupled with changes in cerebral blood flow in the ipsilateral thalamus out to 3 months post-stroke (46). However, to our knowledge, this is the first-time changes in cerebrovascular remodelling have been investigated in the ipsilateral hippocampus at a chronic phase post-stroke. Together, these results suggest that post-stroke angiogenesis and/or vasculogenesis may be a common process in remote brain regions implicated in SND, including the hippocampus. Although the specific underlying mechanisms of the remote angiogenesis were beyond the scope of the study, previous studies in the hippocampus have reported early expression changes in the VEGF/VEGFR pathway which precedes neovascularisation after cerebral ischemia (38). Angiogenesis arising from the subventricular zone has been shown to play a role in brain remodelling after 2h MCAo in rats (47). Within the hippocampus the DG contains neural progenitor cells, which are a source of trophic factors, similar to the subventricular zone. Therefore, it is not unreasonable to extrapolate that similar processes are occurring within the hippocampus contributing to the angiogenesis we observed in this study. However further confirmatory investigation is required.

When investigating vessel diameter, we did not observe any changes out to 28 days post-stroke in any region. In contrast, previous studies found that the vessels present in the peri-infarct area were larger diameter on average to at least 28 days post-stroke (40, 48, 49). This discrepancy may be explained by previous studies which used a protocol that measured vessel diameter of perfused fluorescein isothiocyanate labelled gelatin/albumin, whereas, we measured the vessel diameter of lectin labelled fixed brain sections for this study. We did, however, observe a decrease in average vessel diameter at 84 days in the peri-infarct area and CA1. To the best of our knowledge, this is the first ever report of cerebrovascular narrowing in regions at a distance to the primary infarction in the recovery phase of stroke. Amyloid-β oligomers are known to interfere with vascular function (13), and we recently reported deposition of amyloid-β around cerebral vessels at 84 days post-stroke in these animals (11). Interestingly, we demonstrated that vessels with amyloid-β deposited around their walls were narrower than those without amyloid-β accumulation. Amyloid-β can affect vasomotor regulation by enhancing penetrating arteriole constriction and diminishing vessel dilatation (50). Further, cerebral capillary narrowing by pericytes has been reported previously following vessel recanalization in the hyper-acute phase of large vessel stroke (51) and in response to amyloid-β oligomers in Alzheimer’s disease (52). Such reductions in vessel diameter are likely to have profound effects on reducing blood flow, especially in capillary sized vessels, where the effects of reduced diameter on capillary flow are not accurately estimated by Poiseuille’s law and exceed the r^4^ effect of vessel radius on flow (53, 54). As we reported previously, persistent neuronal loss was observed in the CA1 out to 84 days post-stroke, along with deficits in hippocampal-dependent tasks (Figure 6 (11)). Based on the present findings, we considered that the delayed reduction of vessel diameter mediated by amyloid-β oligomers in the CA1 subregion of the hippocampus may partly contribute to ongoing cognitive impairment after cortical stroke.

Vascular leakage was increased in the peri-infarct area and ipsilateral hippocampus as measured by IgG extravasation. The IgG labelling suggests that the competence of the BBB in the peri-infarct and hippocampus was compromised. Similar findings out to 1 month post stroke have been previously reported in the peri-infarct region by us and others using IgG (55) and Evans blue (56) extravasation, as well as high molecular weight dextran (41). BBB disruption has also been observed at different post-stroke timepoints in regions remote from the infarct, including the hippocampus (56) and contralateral hemisphere (57). Here, we showed that this leakage persists out to 84 days post-stroke in the peri-infarct area and CA1 subregion of the ipsilateral hippocampus. This may have contributed to the SND and the persistent cognitive deficits observed in these animals as reported previously (Figure 6, see (11)). While we documented BBB leakage in peri-infarct area and ipsilateral hippocampus, the mechanisms of BBB dysregulation appeared to be brain region-specific. We hypothesise that BBB leakage observed in the peri-infarct area is linked with the observed concomitant increase in MMP9 expression early post-stroke and then the decrease in tight junction protein Claudin 5 longer term. Interestingly as mentioned above, at 84 days post-stroke we observed the build-up of amyloid-β within the vasculature. Increased levels of amyloid-β oligomers have been shown to influence endothelial permeability and the expression of tight junctions *in vitro* (58). Further investigation is required to determine the specific cause of the protein changes observed here, including tight junction markers, MMPs and tissue inhibitors of metalloproteinases. Unlike the peri-infarct area, there was no difference in either Claudin 5 or MMP9 observed at 84 days in the hippocampus, suggesting a different underlying mechanism. In the hippocampus, we observed a significant decrease in the PDGFRβ+ pericyte coverage of the vasculature at 7 and 84 days post-stroke in the CA1 subregion and at only 84 days in the DG subregion of the hippocampus compared with sham animals. Pericytes have an important function in the maturity of blood vessels and the integrity of the BBB (43, 59). We hypothesise that the lack of pericyte coverage on the newly formed vessels in the hippocampus may be a contributing factor to the BBB leakage observed in this study. Our hypothesis is supported by Zbesko et al (2018) who showed similar findings at 7 weeks post-stroke within the infarct core and peri-infarct areas (60). A key future experiment would be to identify these brain region-specific BBB dysregulation mechanisms to develop targeted therapies for mitigating the progression of hippocampal BBB leakage.

In conclusion, we demonstrated that cortical photothrombotic stroke causes cerebrovascular changes that are long term and persistent in the CA1 sub-region of the hippocampus, an area critical in cognition. Indeed, we previously demonstrated persistent cognitive impairment and provided critical insights into the underlying mechanisms of SND in the hippocampus in these animals (see Figure 6). Interestingly, we showed associations between amyloid-β and reduced vessel diameter, which likely contributed to the persistent neuronal loss and cognitive impairment observed in these animals (11). Further exploration into the specific mechanisms underlying post-stroke SND and cognitive impairment is required, however our findings indicate that hippocampal vasculature and BBB function may represent an important therapeutic target to mitigate the progression of post-stroke SND and cognitive impairment.

## Supporting information

Supplemental Methods and Results

## Author Contributions

Conceptualization, S.S.-B., F.R.W., M.N., L.K.O.; Data curation, S.S.-B., R.J.H and L.K.O.; Formal analysis, R.J.H, S.S.-B., R.R. and L.K.O.; Funding acquisition, F.R.W., M.N., and L.K.O.; Methodology, S.S.-B., R.J.H and L.K.O.; Supervision, F.R.W., M.N. and L.K.O.; Writing–original draft, R.J.H, D.J.B. and L.K.O.; Writing–review and editing, R.J.H, S.S.-B., D.J.B, R.R., R.J.T., S.M.S., L.E.C.-P., F.R.W., M.N. and L.K.O. All authors have read and agree to the published version of the manuscript.

## Declaration of conflicting interests

The author(s) declare no potential conflicts of interest with respect to the research, authorship, and/or publication of this article.

### Acknowledgements

LKO, FRW and MN acknowledge ongoing support from NHMRC Centre for Research Excellence in Stroke Recovery and Rehabilitation. LKO and SSB acknowledge support from Research Advantage for ECR Higher Degree by Research (HDR) Scholarship and Greaves’ Family Postgraduate Scholarship in Medical Research (HMRI 1054). LKO acknowledges support from the University of Southern Queensland Research Capacity Building Grant, the International Society for Neurochemistry (ISN) Career Development Grant, IBRO-APRC Travel & Short Stay Grant, Hunter Medical Research Institute (HMRI 896) and The University of Newcastle, Australia. RJT and LECP acknowledge support from NeuroSurgical Research Foundation and Perpetual.

