## Supplemental Methods and Results for "Leakage beyond the primary infarction: A temporal analysis of cerebrovascular dysregulation at sites of hippocampal secondary neurodegeneration following cortical photothrombotic stroke"

### Supplementary Methods

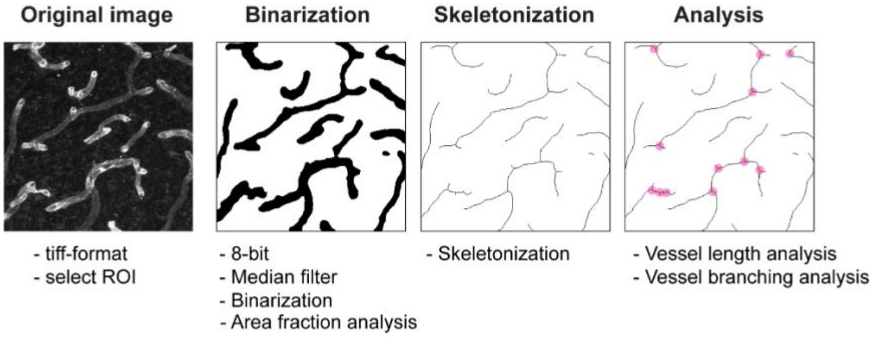


**Supplementary Figure 1:** Illustrates the process of digital modelling based on previously established automated ImageJ (Fiji) script. This process generated the formation of thresholded and skeletonized images, which were then used for morphological analysis.

### Supplementary Results - Raw Western Blots

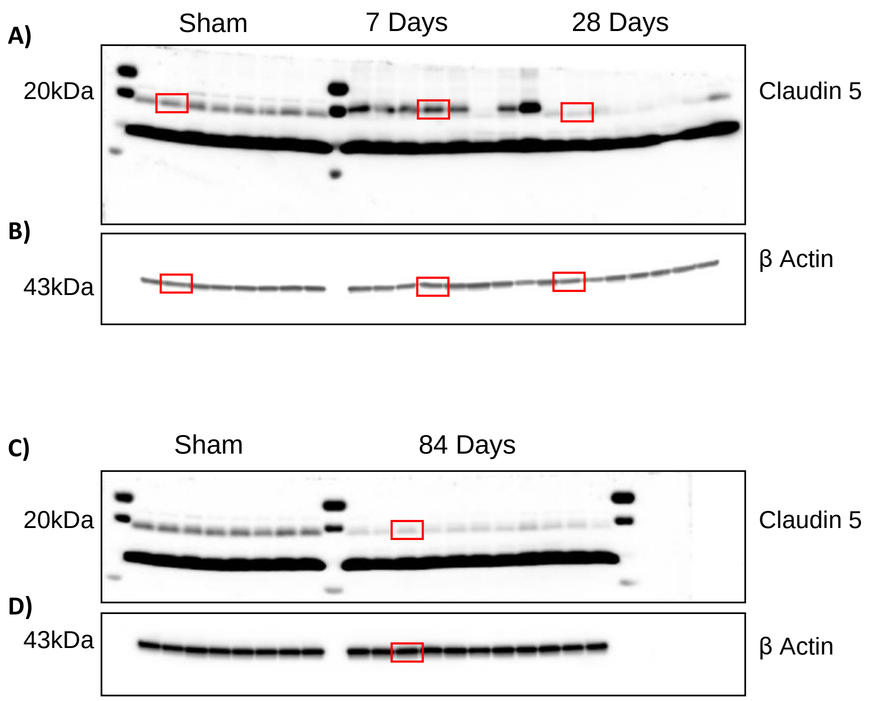


**Supplementary Figure 2: Full unedited blots for Figure 5A – used for the analysis of Claudin 5 in the peri-infarct area.** The panels are chemiluminescent images taken using an Amersham Imager 600, which provides information of the molecular weight/size of the bands (weights depicted to left of blots). The red boxes indicate the bands featured in Figure 5A. The signals of the bands from the original, unprocessed immunoblots were measured using Amersham Imager 600 Analysis Software.

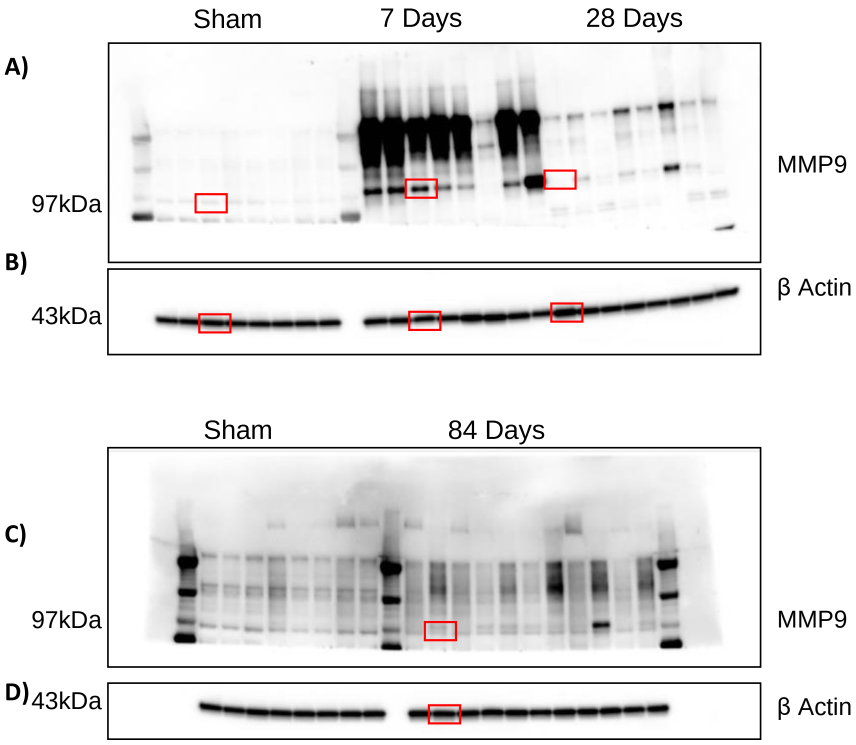


**Supplementary Figure 3: Full unedited blots for Figure 5B – used for the analysis of Matrix metalloproteinase 9 (MMP9) in the peri-infarct area.** The panels are chemiluminescent images taken using an Amersham Imager 600, which provides information of the molecular weight/size of the bands (weights depicted to left of blots). The red boxes indicate the bands featured in Figure 5B. The signals of the bands from the original, unprocessed immunoblots were measured using Amersham Imager 600 Analysis Software.

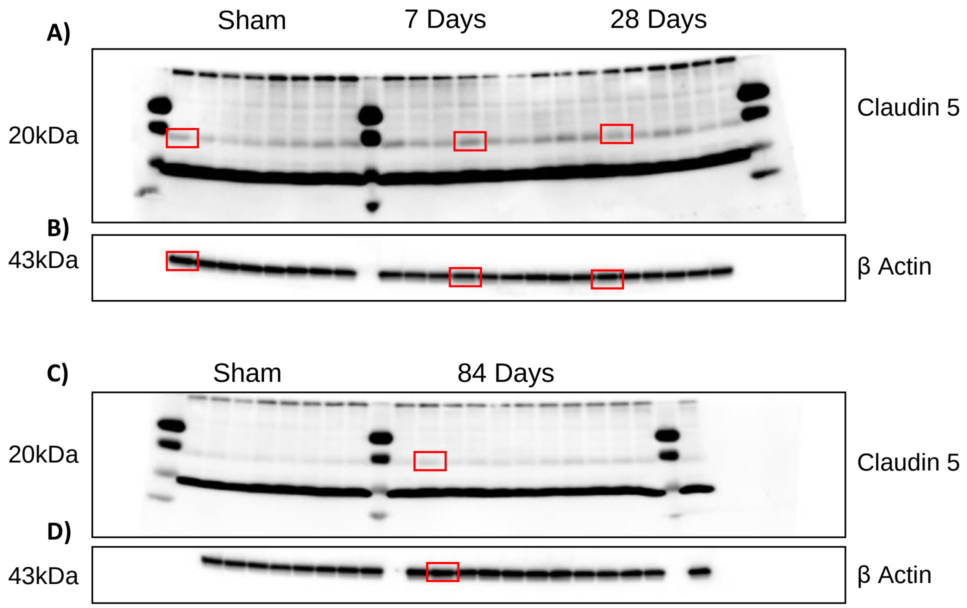


**Supplementary Figure 4: Full unedited blots for Figure 5C – used for the analysis of Claudin 5 in the hippocampus.** The panels are chemiluminescent images taken using an Amersham Imager 600, which provides information of the molecular weight/size of the bands (weights depicted to left of blots). The red boxes indicate the bands featured in Figure 5A. The signals of the bands from the original, unprocessed immunoblots were measured using Amersham Imager 600 Analysis Software.

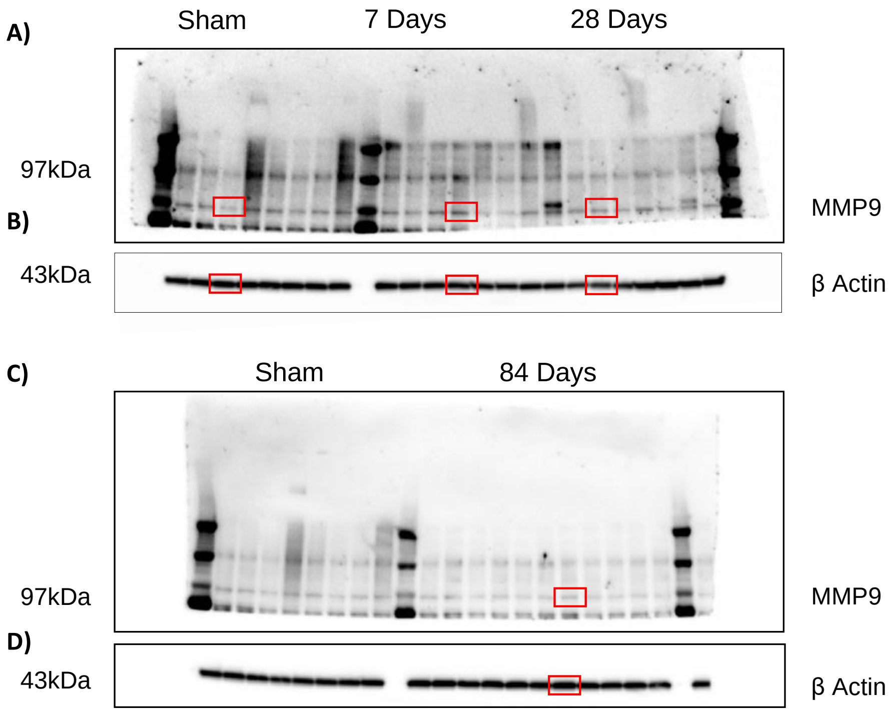


**Supplementary Figure 5: Full unedited blots for Figure 5D – used for the analysis of Matrix metalloproteinase 9 (MMP9) in the hippocampus.** The panels are chemiluminescent images taken using an Amersham Imager 600, which provides information of the molecular weight/size of the bands (weights depicted to left of blots). The red boxes indicate the bands featured in Figure 5D. The signals of the bands from the original, unprocessed immunoblots were measured using Amersham Imager 600 Analysis Software.
